# Ecological habitat partitioning and feeding specialisations of coastal minke whales (*Balaenoptera acutorostrata*) using a designated MPA in northeast Scotland

**DOI:** 10.1101/2021.01.25.428066

**Authors:** Kevin P. Robinson, Connor C.G. Bamford, William J. Brown, Ross M. Culloch, Ciaran J. Dolan, Rebecca Hall, Grace Russell, Theofilos Sidiropoulos, Evgenia Spinou, Texa M.C. Sim, Elice Stroud, Genevieve Williams, Gary N. Haskins

## Abstract

In the design of protected areas for cetaceans, spatial maps rarely take account of the life-history and behaviour of protected species relevant to their spatial ambit, which may be important when modelling population trends or assessing susceptibility to anthropogenic threats. In the present study, we examined the distribution and feeding behaviours of minke whales by age-class (adults *versus* juveniles) from long-term studies in the Moray Firth in northeast Scotland, where a Marine Protected Area (MPA) has recently been designated. Data were collected from dedicated boat surveys between 2000 and 2019, during which 657 encounters with 774 whales of confirmed age-class (444 juveniles and 330 adults) were recorded from 50,041 km of survey effort, resulting in 224 individual follows. Feeding/foraging whales were documented in 84% of the encounters. Adults and juveniles were occasionally seen together, but their distributions were not statistically correlated, and GIS revealed spatial separation by age-class―with juveniles preferring shallow, inshore waters with sandy-gravel sediments and adults preferring deeper, offshore waters with steep benthic slope. Whilst adult minkes employed a range of “active” prey-entrapment specialisations, showing seasonal flexibility in their targeted prey with interindividual variation, juveniles almost exclusively used “passive” (low energy) feeding methods, targeting low-density patches of inshore prey. These findings corroborate the need to incorporate demographic/behavioural data into spatial models when identifying priority areas for protected cetaceans and may be important to adaptive management objectives for the species in the Moray Firth MPA.

## Introduction

The minke whale (*Balaenoptera acutorostrata* Lacépède) is the smallest and most abundant of the baleen whales in UK waters. Approximately 9,000 occur in the North Sea [1], with most sightings in the northern North Sea and primarily inshore, in shelf waters less than 200 metres deep [2]. The highly productive waters of the Moray Firth in northeast Scotland (57° 41′ N, 2° 40′ W) attract above-average densities of minke whales relative to adjacent and wider Scottish waters [3], affording rich feeding grounds for the species during the summer and autumnal months [2,4]. Accordingly, the Southern Trench in the outer Moray Firth (Fig 1) was recently appointed as a Marine Protected Area (MPA) [5] for the protection of these whales in this coastal northeast location.

**Fig 1.**
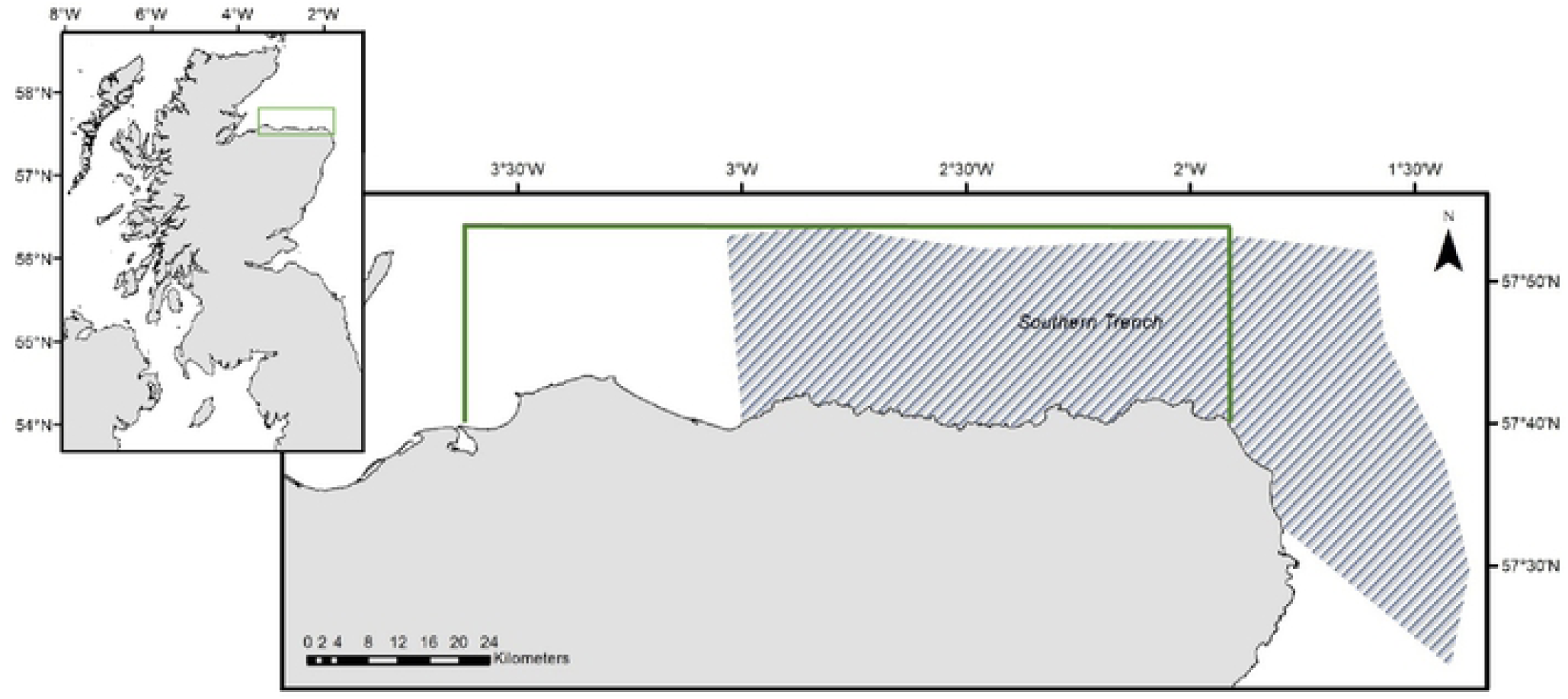
The position of the 1,980 km^2^ study area (green border) and the boundaries of the Southern Trench MPA (shaded) along the southern coastline of the outer Moray Firth in northeast Scotland in the northern North Sea.

The mapping of high-density areas from distributional sightings data is a crucial first step in the design of protected area management for cetaceans [6]. However, spatial maps rarely take account of the life-history and behaviour of protected species relevant to their spatial ambit [7] which may be important for conservation management. Robinson et al. [2] note the high percentage of juvenile minkes (comprising ∼60% of all sightings) frequenting the Moray Firth, and their nearshore predilection for sandeel (*Ammodytes marinus*) predicted habitat. Whilst the low energetic cost of swimming in these whales allows them to exploit environmental conditions over large spatial scales [8-9], juvenile minkes may be less efficient foragers than adults and are potentially displaced from optimal feeding areas by their larger conspecifics, forcing them to forage for alternate resources [10]. Moreover, individual whales, and experienced adults in particular, may further employ unique feeding specialisations for entrapping their prey [11], resulting in intrapopulation variation in resource use and dietary plasticity in the species.

Certain eco-geographic variables (EGVs) in the marine environment favour predator and prey species alike [12-13], and factors such as ocean floor topography, water depth, sea bottom sediment, tidal fronts and water temperature may exert a strong influence on the distribution of minke whales throughout their range, at both fine- and meso-scale levels [2,14-18]. In the following study, we investigated whether whales of different age-classes exhibited differences in their spatial occurrence and habitat use in the coastal Moray Firth. The spatial distribution of adult *versus* juvenile whales was subsequently examined with respect to the proximity of recorded sightings to shore and the physiographic predictors water depth, benthic slope and sediment-type. Observational data were further analysed to investigate feeding methods and dietary preferences in the species, as well as intraspecific variations in the feeding strategies employed by individual specialists. The primary focus of this investigation was to identify priority habitats within the designated MPA and surrounding area which might be biologically important for the species and the susceptibility of discrete demographic subgroups to contiguous anthropogenic activities.

## Materials and methods

### Data collection

Sightings data were collected during dedicated boat-based surveys within a 1,980 km^2^ area of the southern Moray Firth between May and October 2000 to 2019 (Fig 1). The surveys were carried out using rigid inflatable boats with a crew of at least two experienced and up to six additional trained observers searching for whales using a continuous scanning method, after Mann [19]. Only the initial sighting for each whale was used in the following investigation to avoid data replication/autocorrelation. After each encounter, the search effort was further directed to previously un-surveyed areas, to minimise repeated encounters of the same individual whales and to maximise spatial coverage during boat surveys.

Cues used to locate whales during surveys included the presence of feeding birds [4] in addition to direct observations of the animals themselves when travelling or surface feeding [2]. When a sighting was made, the time, immediate geographic position of the animal(s) (corrected for distance), behaviour (feeding/foraging or travelling) and age-class (adult/juvenile) of the whale(s) were recorded where possible. Adult minkes were defined as large, dark coloured animals >6.5 metres in length, whilst juveniles were defined as lighter, olive-coloured animals <6.5 metres, after Mitchell E, Kozicki [20]. Sightings which could not be assigned to an age-class, due to the briefness of the encounter, evasiveness of the animal or poor lighting conditions, for example, were not included in the following analysis (*n* = 186).

An ethogram was used to describe the surface feeding specialisations used by individual whales during observed predation events (Table 1). A minimum of six spotters were tasked with tracking each whale during individual focal-follows, providing 360° visual coverage around the survey vessel so ensure that no surfaces / behaviours were missed. All whale follows were conducted off-survey effort (termed encounter effort), with boat distances maintained between 50 and 300 metres from the subject during sampling periods of up to 30 minutes. A medium-mesh, extendable landing net (Aquascape Ltd, UK) was also used for the *ad-libitum* recovery of prey species for species identification, and length measurements were recorded *in situ* before sampled prey were returned back to the sea.

**Table 1.** Ethogram detailing the surface feeding behaviours / prey entrapment methods employed by minke whales frequenting the coastal Moray Firth.

| Behaviour | Description |
| --- | --- |
| *General* |  |
| Passive (bird-associated) feeding | Whales exploit concentrations of baitfish compacted together at the water's surface by flocks of feeding seabirds from above and schooling predatory fish from below |
| Active feeding | Whales actively corral the baitfish themselves, showing multidirectional surfacing followed by a variety of feeding strikes (aerial, surface or sub-surface lunges) in dorsal, ventral or lateral planes |
| *Active Feeding* |  |
| <u>Pre-strike</u> |  |
| Head slap | Whale lifts its head above the water and audibly slaps its chin down on the surface |
| Depth-charge | After surfacing and re-submersing, the whale exhales air forcefully underwater creating a cacophony of bubbles at the surface |
| <u>Strike</u> |  |
| Sub-surface lunge | Whale strikes below the surface producing a wave of white water, but the body is not seen |
| Surface lunge | Whale breaks the surface of the water and the arcing body is visible above the surface |
| Aerial lunge | Whale lunges high out of the water and the entire head and body is viewed |
| Plane of strike | The whale strikes the prey in either dorsal, ventral or lateral (either right or left side down) planes |
| Speed of strike | The strike is either fast and powerful or slow and progressive |
| <u>Post-strike</u> |  |
| Roll | The whale rolls to the right- or left-hand side post-striking |

### Spatial analysis

A rectangular grid of the geographic study area was created using ArcGIS Desktop 10.6.1 (ESRI, USA), with each grid cell measuring 0.25 km^2^. The complete survey data from 2000 to 2019, consisting of 50,041 km of on-effort track data and 774-point locations of minke whales, were subsequently imported into ArcGIS to examine the spatial distribution of adult and juvenile minke whales with respect to the underlying EGVs sediment type, water depth, bathymetric slope and also the proximity to shore. The sediment data were provided under licence from the British Geological Survey and depth data were obtained from GEBCO (30-arc second dataset) [21]. The slope layer was derived from the depth data using a custom GIS workflow, whilst proximities of sightings to the shore were calculated using a geodesic Euclidean Distance tool. After a successive processe of simplification and classification, all layers were converted to Boolean maps for generation of the respective values within each 0.25 km^2^ grid cell. Moran’s *I*-tests were run using the ‘ape’ package in R 3.1.2 (http://www.r-project.org) [22] to test for the presence of spatial autocorrelation in the whale sightings per grid cell for each survey year.

### Data modelling

Generalised additive models (GAMs) were used to examine the non-linear relationship between adult and juvenile minke whales and their habitat for all sightings data from 2000 to 2019 inclusive. The data were modelled with R 3.1.2 using logistic regression with a binomial response for the presence/absence of adult and juvenile whales per grid cell with smoother terms derived from penalised regression splines using the *mgcv* package in R [23]. Since presence is a probabilistic function mainly affected by species abundance and detectability [24], assuming that the detectability of whales across all habitats was constant, absences were subsequently associated with habitats in which abundance was low. The explanatory model terms (water depth, slope and proximity to shore) were treated as continuous variables, with spline smoothers initially fitted to each term in the model. Selection of significant terms was subsequently carried out using a backward selection method. Models with the lowest cross-validation scores were selected [22] and outputs examined for patterns of residuals to validate the models using the Akaike Information Criteria (AIC) statistic for final model selection.

## Results

The spatial distribution of all minke whale sightings of known age-class in the study area is shown in Fig 2. Between 2000 and 2019, a total of 774 individuals were recorded from 657 encounters. Feeding/foraging whales were predominantly encountered over travelling/resting whales (84% *versus* 16%), whilst juvenile animals were more frequently encountered than adults (444 juveniles c.f. 330 adults). Results from the Moran’s *I*-tests (p-values) revealed no autocorrelation in the sightings data for any of the survey years examined, and pairwise comparisons of sightings per grid cell revealed that the spatial sightings of adults and juveniles were not statistically correlated. In addition, GIS resolutions inferred a strong association by juvenile minkes for shallow (< 50 m deep), inshore waters (mean distance from coast = 2.79 km), with low benthic topography, whereas adult whales were more typically associated with deeper waters (between 20 and 80 m), further from the coast (mean distance = 5.39 km), over areas of steeper benthic slope (Figs 3a to c, Table 2). Sightings of juveniles were also strongly correlated with sandy gravel sediment (Spearman’s Rank Correlation r^2^ = 0.86, p < 0.001, *n =* 444), whilst adults were predominantly associated with areas of muddy sand (Spearman’s Rank Correlation r^2^ = 0.56, p < 0.05, *n =* 330) (Fig 3d).

**Table 2.** The mean water depth, slope and proximity to shore of adult and juvenile minke whale sightings recorded in the outer Moray Firth from 2000 to 2019.

| Adults ($n = 330$) | Mean ± SD | Min | Max |
| --- | --- | --- | --- |
| *Depth (metres)* | $50.9 \pm 40.8$ | 9.2 | 207.0 |
| *Slope (degrees)* | $2.20 \pm 1.42$ | 0.03 | 12.20 |
| *Proximity to shore (km)* | 5.39 | 1.06 | 23.02 |
| Juveniles ($n = 448$) | Mean ± SD | Min | Max |
| *Depth (metres)* | $28.2 \pm 52.3$ | 4.5 | 202.6 |
| *Slope (degrees)* | $0.90 \pm 0.99$ | 0.02 | 7.77 |
| *Proximity to shore (km)* | 2.79 | 0.92 | 20.92 |

**Fig 2.**
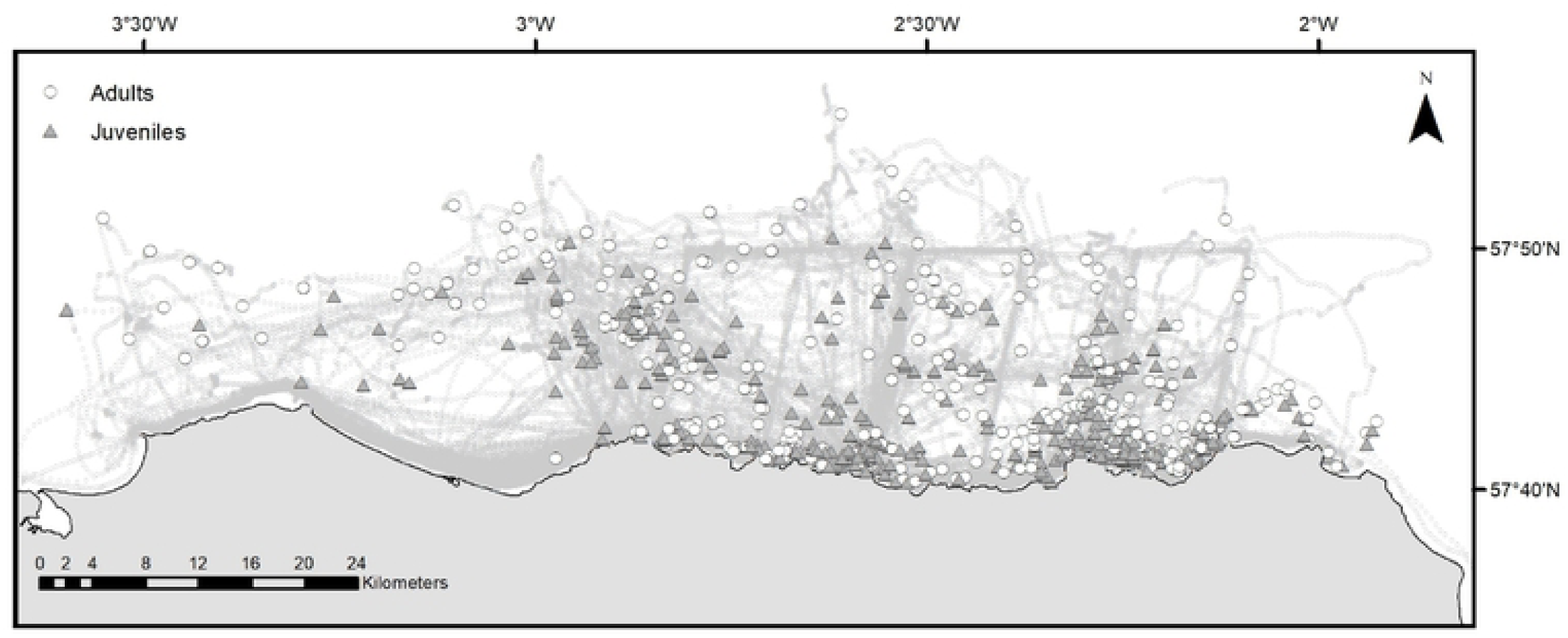
The spatial distribution of adult and juvenile minke whale *Balaenoptera acutorostrata* sightings recorded in the Moray Firth study area between May and October 2000 to 2019. A total of 50,041 km of boat survey effort was conducted resulting in 774 sightings confirmed to age-class.

**Fig 3.**
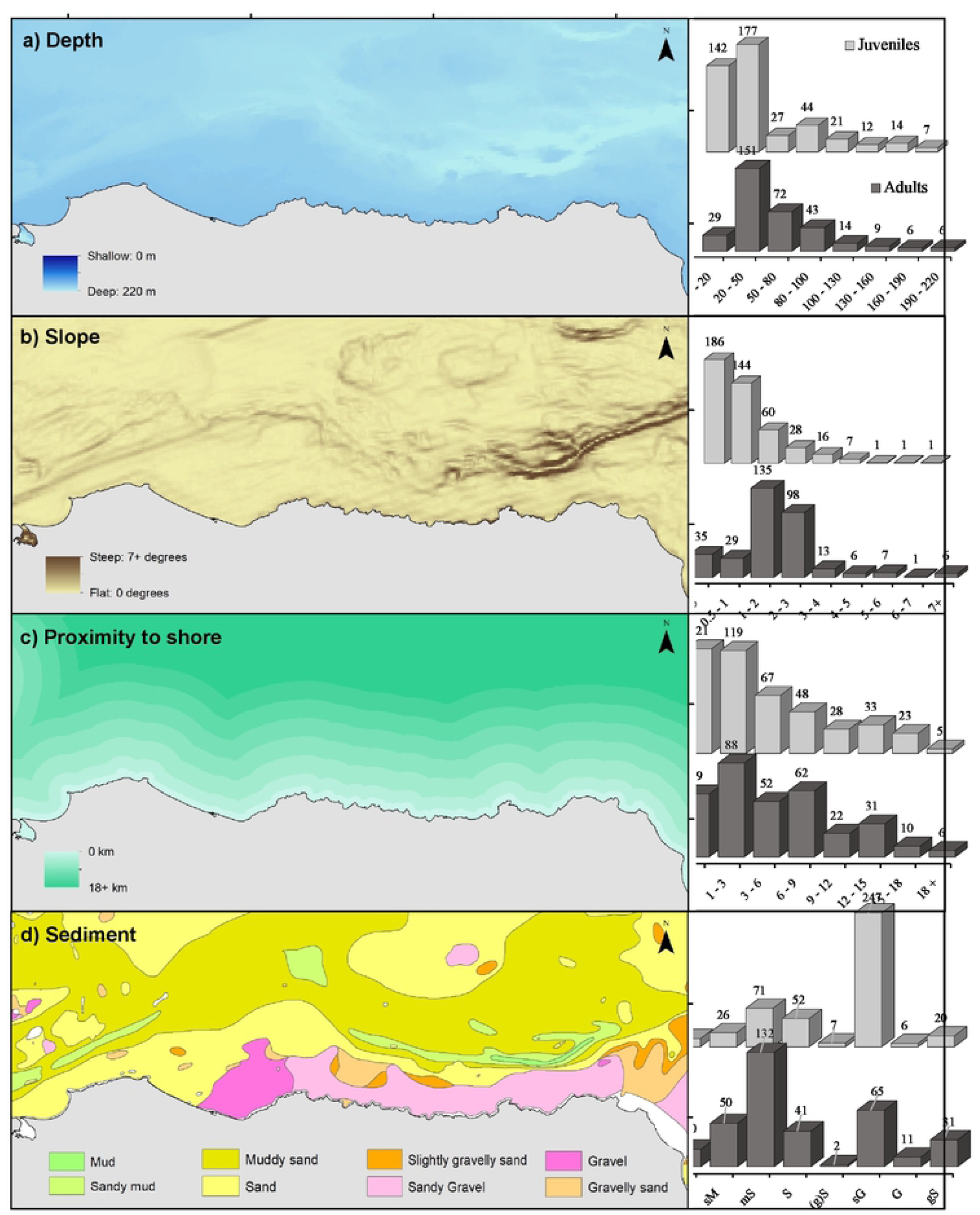
The spatial associations of adult and juvenile minke whales in the Moray Firth study area with respect to the eco-geographic variables (a) water depth, (b) benthic slope, (c) proximity to shore and (d) sea bottom sediment.

The GAM results showed that slope had a positive, non-linear effect (p < 0.01) on the distribution of adult whales, but was not significant for juveniles (Table 3). Conversely, distance from shore and water depth were found to be important predictors for the distributions of both adults and juveniles alike (Table 3). The final GAM for adult whales explained 64% of the deviance in the form: *n* ∼ *s* (slope) + *s* (depth) + *s* (proximity to shore), where *n* represents the sightings rate (no. of sightings per km) and *s* the smoother function of each covariant. The final GAM for juvenile whales explained 36% of the deviance in the form: *n* ∼ *s* (depth) + *s* (proximity to shore).

**Table 3.** Generalised additive model (GAM) results for non-linear minke whale relationships with eco-geographic variables as determined from the number of whales per 0.25 km^2^ grid cell surveyed. Significant p-values are shown in bold, ns = not significant.

| Predictor<br>(age- class) | Water depth |  |  | Slope |  |  | Sediment type |  |  | Proximity to shore |  |  |
| --- | --- | --- | --- | --- | --- | --- | --- | --- | --- | --- | --- | --- |
|  | edf | *F* | p | edf | *F* | p | edf | *F* | p | edf | *F* | p |
| Adults (*n* = 330) | 1 | 7.62 | < **0.01** | 1 | 16.54 | < **0.001** | NA | NA | ns | 1.322 | 2.92 | < **0.05** |
| Juveniles (*n* = 448) | 1 | 5.69 | < **0.01** | 1 | 0.02 | ns | NA | NA | < **0.001** | 1.086 | 3.69 | < **0.05** |

283 feeding strikes were recorded from 224 follows of individual minke whales, during which “passive” and “active” feeding behaviours were documented (Table 4). Active feeding was widely recorded in adults (Fig 4) but was rarely observed in juveniles (just 4% of the behaviours), as juveniles almost exclusively used passive (bird-associated) feeding methods instead (Table 4). Whales of both age-classes were recorded engulfing prey using lateral and/or dorsal planes when striking. Lateral strikes were chiefly orientated right-side down or with the whale rolling to the right post-strike, with just 10% of the animals performing left-sided manoeuvres (Table 4). Behaviours such as head slapping and depth-charging (blow under water after diving) were only ever used by adult specialists, and aerial lunges (where the whale exited the water when feeding) were only performed in the absence of surface feeding seabirds. A total of 47 recognisable whales were recaptured during the study period on two or more separate occasions. Of those individuals recaptured during different months in the same year (*n* = 11) or during different survey years (*n* = 14), on each occasion the same specific prey entrapment methods (orientation/type) were observed.

**Table 4.** Surface feeding specialisations recorded during individual focal follows (*n* = 224) of minke whales in the Moray Firth between 2000 and 2019 inclusive.

|  | Adults | Juveniles | All |
| --- | --- | --- | --- |
| *Feeding type* |  |  |  |
| Passive (bird-associated) feeding | 16 | 160 | 176 |
| Active feeding / corralling | 101 | 6 | 107 |
| TOTAL | 117 | 166 | 283 |
| *Pre-Strike Activity* |  |  |  |
| Head slap | 9 | 0 | 9 |
| Depth-charge | 21 | 0 | 21 |
| *Type of Strike* |  |  |  |
| Sub-surface | 36 | 132 | 168 |
| Surface | 67 | 29 | 96 |
| Aerial | 14 | 5 | 19 |
| *Plane of Strike* |  |  |  |
| Dorsal | 19 | 61 | 80 |
| Ventral | 7 | 0 | 7 |
| Lateral | 91 | 105 | 196 |
| *Right side down* | 83 | 95 | 178 |
| *Left side down* | 8 | 10 | 18 |
| *Speed of Strike* |  |  |  |
| Fast/powerful | 99 | 130 | 229 |
| Slow/progressive | 18 | 36 | 54 |
| *Post-Strike Roll* |  |  |  |
| On the right-hand side | 95 | 13 | 108 |
| On the left-hand side | 9 | 0 | 9 |

**Fig 4.**
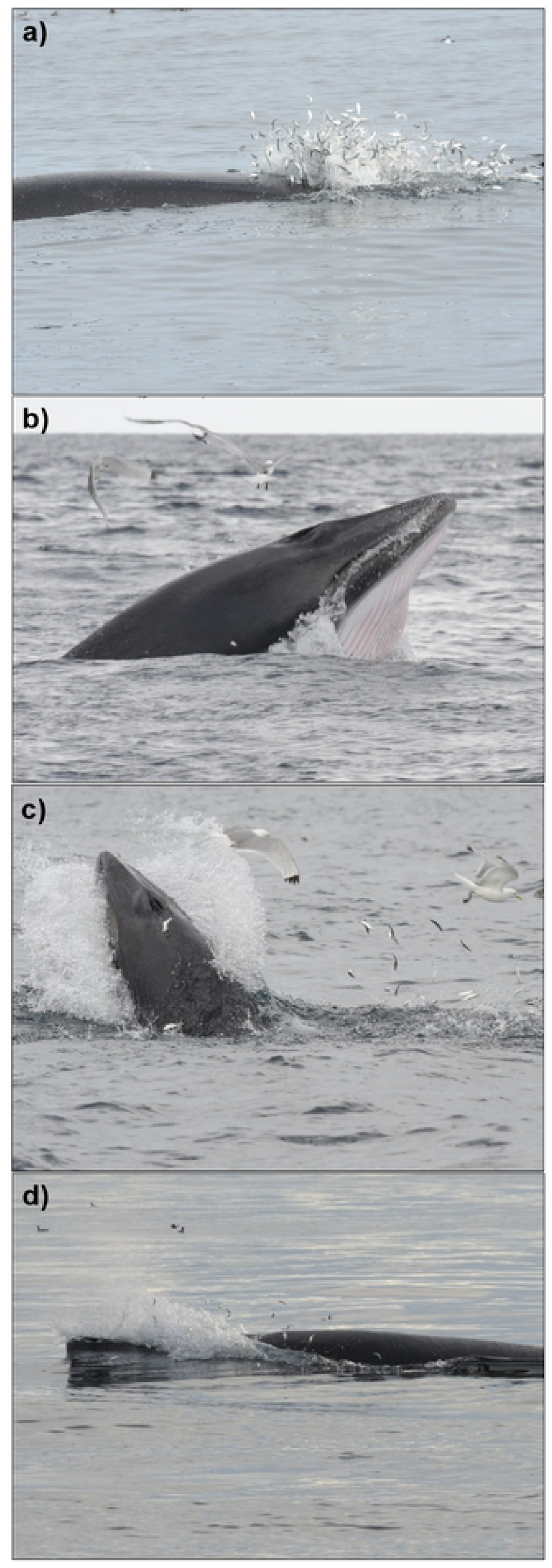
“Active” adult minke predation events upon (a and b) juvenile herring and (c and d) pre-wintering sprat. Photographs: Kevin Robinson.

Prey items were recovered from 95 feeding events between 2002 and 2017 and identified prey species included lesser sandeels (*A. marinus*), herring (*Clupea harengus*) and sprat (*Sprattus sprattus*) (Table 5). Juvenile whales primarily targeted year 0-1 sandeels (measuring 86 to 118 mm in length) [25]. However, larger prey items, including year 0-3 sandeels, juvenile herring and pre-wintering sprat, were consistently recovered from adult feeding events (Table 5). Sandeels were targeted by adults and juveniles alike during all study months, May to October inclusive. However, juvenile herring were preferentially targeted by adults from early July, whilst sprat were targeted by both adults and juveniles alike from late August to October. The recorded seasonal changes in the proximities of animals to the shore, the feeding methods employed and the prey species sampled for each age-class are summarised in Table 6.

**Table 5.** Fish prey species sampled from individual feeding events (n = 95) of adult and juvenile minke whales between 2002 and 2017.

| Age Class | Sandeel |  |  | Herring |  |  | Sprat |  |  |
| --- | --- | --- | --- | --- | --- | --- | --- | --- | --- |
|  | Mean length (mm) | Range (mm) | *n* | Mean Length (mm) | Range (mm) | *n* | Mean Length (mm) | Range (mm) | *n* |
| Adults (*n* = 52) | 119 | 85 - 163 | 14 | 220 | 196 - 321 | 22 | 122 | 92 - 156 | 16 |
| Juveniles (*n* = 43) | 104 | 86 - 118 | 41 | 0 | 0 | 0 | 107 | 94 - 122 | 2 |

**Table 6.** Observed intra-seasonal changes in minke whale feeding characteristics from the pooled dataset 2000 to 2019.

| Interval | Age-class | Feeding methods |  | Prey species |  |  | Proximity to shore |  |  |
| --- | --- | --- | --- | --- | --- | --- | --- | --- | --- |
|  |  | Passive | Active | Sandeel | Herring | Sprat | 0-5 km | 5-10 km | >10 km |
| May to Jun | Adults | 9 | 30 | 10 | 1 | 0 | 5 (10.4%) | 29 (60.4%) | 14 (29.2%) |
|  | Juveniles | 74 | 0 | 28 | 0 | 0 | 85 (68.5%) | 36 (29%) | 3 (2.5%) |
| Jul to Aug | Adults | 5 | 59 | 2 | 17 | 3 | 44 (23.4%) | 126 (67%) | 18 (9.6%) |
|  | Juveniles | 83 | 2 | 11 | 0 | 0 | 136 (47.5%) | 142 (49.7%) | 8 (2.8%) |
| Sep to Oct | Adults | 2 | 12 | 2 | 4 | 13 | 11 (11.7%) | 59 (62.8%) | 24 (25.5%) |
|  | Juveniles | 3 | 4 | 2 | 0 | 2 | 16 (47.1%) | 17 (50%) | 1 (2.9%) |

## Discussion

Occurrences of baleen whales on their feeding grounds are typically linked to environmental variables which influence the distribution of their prey [26-27]. In the present study, benthic slope, water depth and proximity to shore were found to be significant predictors for the occurrence of adult minke whales, whilst proximity to shore, water depth and sediment-type were the most important predictors for juveniles. Juvenile whales were also found to be more prevalent than adults within the Moray Firth study area, representing approximately 60% of all sightings determined to age-class. Inevitably, juveniles utilising the study area may have been resighted on separate survey days during the same year, and this may have biased this finding. However, since juvenile animals were typically unidentifiable from natural markings (i.e. dorsal fin nicks / tears) such as those more reliably observed in adults, it was not possible to correct for this bias. Juvenile and adult minkes were occasionally seen foraging together, however their distributions were negatively correlated, suggesting intrapopulation partitioning by age-class in the species. Haug et al. [28] reported that during their northward migration minke whales show segregations by sex and size, with adult females and juveniles inhabiting more coastal areas and adult males tending to remain further offshore. In ecological systems, age- or sex-based differences may arise as a necessary consequence of body-size or development such that partitioning might occur as a by-product of ontogeny [e.g. 10].

Certainly, identified adults using the Moray Firth were seen to target larger prey than their juvenile counterparts. Juveniles almost exclusively targeted year 0-1 sandeels in the study area, as confirmed from the sampling of targeted prey during feeding events, but adults showed a seasonal flexibility switching between sandeels (year 0-3), herring and sprat (Fig 4)―the three species contributing between them up to 86% of the total fish biomass of the Moray Firth [29]. Sandeels are a short-lived, benthic fish, strongly associated with sandy-gravel sediments [30], to which the juvenile whales were also closely correlated in this study. Conversely, herring and sprat are mid-water, shoaling species that occur in deeper, shelf waters [31-32] where adult whales were more typically encountered. From June to August, juvenile herring seemed to be preferentially targeted by adult whales over sandeels, but from August to October pre-wintering sprat were targeted over herring, indicating prey-switching by the species. Perhaps the complex schooling behaviour and strong predator avoidance shown by herring [32] reduces the whales’ preference for herring when sprat are more widely accessible. However, predators naturally show heritable flexibility in their resource preferences when options are limited, or when an alternative, high-valued resource becomes more widely abundant [33]. Within the Moray Firth study area in 2006, for example, following the EU-wide ban on the North Sea sandeel fishery [34], disproportionate numbers of both adult and juvenile minkes were sighted inshore, visibly profiting from high densities of sandeel prey (K Robinson pers. observation). It is widely reported that minke whales respond to seasonal changes in the abundance of their prey [e.g. 15,28], and this is assumed to occur when prey densities surpass a particular threshold that is energetically profitable to foraging whales for switching to occur [2,9]. This could conceivably explain the high interannual and intra-seasonal variability in resource selection noted in this and other UK studies of the species and the apparent plasticity in diet shown by these coastal balaenopterids [e.g. 9,15,17].

When hunting for their prey, minke whales evidently employ a wide range of feeding strategies [e.g. 4,11,18]. In the present study, these included “passive” (bird-associated) and “active” methods for prey entrapment, as first described by Hoelzel et al. [35]. Both adult and juvenile whales invariably pursued patchy resources within the study area, but the active feeding methods used by adults were rarely, if ever, observed in juveniles. Instead, juveniles typically employed low energy, passive prey entrapment methods, targeting small patches of ephemeral prey close inshore [4]. Conversely, adult whales exhibited a broad range of individual specialisations, using a combination of mechanical, acoustic and visual behaviours to actively corral their prey. Observed acoustic behaviours, thought to be used to intimidate prey into a protective response, included head slapping and depth-charging (a forceful blow under water after diving) techniques that have also been described for the species in Canadian waters by Kuker et al. [11]. Interestingly, known (photo-identified) individuals using the Moray Firth study area utilised the same unique specialisations during repeated encounters in different years (K Robinson pers. observation), perhaps alluding to an individually learned component of foraging, resulting in the wide variety of feeding “styles” observed in this species [e.g. 36]. In addition, the majority of feeding whales using the study area showed a clear preference for laterally orientated feeding strikes, showing a 90:10 right-handed bias similar to the handedness index in humans [37]. A skewed ratio for directional lateral feeding has similarly been reported in humpback whales (*Megaptera novaeangliae*) [38] and blue whales (*Balaenoptera musculus*) [39], with individuals showing consistency in the orientation of their feeding strikes or rolls. One well-marked adult recaptured 6 times in the Moray Firth between 2006 and 2019 was only ever recorded using left-handed feeding manoeuvres, suggesting that basic brain lateralisation may be expressed in the same way in cetaceans as in other vertebrates [e.g. 40]. In this context, individual specialisation by these rorqual whales may yield positive benefits for their conservation by adding to the stability of populations [41] and their evolutionary diversification [42].

Clearly, not all parts of an MPA are of equal value for monitoring [6] and management plans aiming to protect a species by targeting “average” resources pose a significant risk when intrapopulation variation exists due to demographic differences [e.g. 43]. Given the spatial differences observed by age-class, the simplification of generalised habitat preferences universally described for the species―i.e. association with the 50-metre isobath, affinity for sandy-gravel sediments and preference for areas of steep topography [e.g. 2, 15, 17]―may subsequently overlook the niche disparities reported herein. The identification and protection of critical habitat can be notoriously difficult in marine ecosystems, and there is a strong need to integrate behavioural and demographic data into spatial models when identifying priority areas for protected cetacean species [e.g. 44]. The nearshore habitats utilised by juvenile minkes, for example, may harbour greater impacts from anthropogenic activities that may be relevant to adaptive management objectives for the species in the Moray Firth MPA. The present findings add greatly to our understanding of the habitat use and ecology of the minke whale frequenting the northeast UK coast, and the subsequent identification of priority areas that may be important to different demographic groups.

## Acknowledgements

All field work was carried out under licence from NatureScot (formerly Scottish Natural Heritage). The sediment data were kindly supplied by the British Geological Survey (BGS) in Nottingham. Thanks go to the numerous colleagues, interns, volunteers, supporters and funders who have made this long-term study possible. We thank the Born Free Foundation (formerly Care for the Wild International) for their long-standing support, and our co-workers and collaborators from the University of Aberdeen, Scottish Agricultural College, University of St. Andrew’s, Scottish Natural Heritage and Whale & Dolphin Conservation.

